# The combination of the functionalities of feedback circuits is determinant for the number and size of attractors of molecular networks

**DOI:** 10.1101/060608

**Authors:** Eugenio Azpeitia, Stalin Muñoz, Daniel González-Tokman, Mariana Esther Martínez-Sánchez, Nathan Weinstein, Aurélien Naldi, Elena R. Álvarez-Buylla, David A Rosenblueth, Luis Mendoza

**Affiliations:** INRIA project-team Virtual Plants, joint with CIRAD and INRA, Montpellier Cedex 5, France.; Instituto de Investigaciones en Matemáticas Aplicadas y en Sistemas, Universidad Nacional Autónoma de México, Apdo. 20-126, 01000 México, D.F., México.; Investigador Cdras CONACYT, Instituto de Ecología, A. C., Antiguo camino a Coatepec 351, El Haya, 91070 Xalapa, Veracruz, México.; Programa de Doctorado en Ciencias Biomédicas, Universidad Nacional Autónoma de México, México.; Instituto de Ecología, Universidad Nacional Autónoma de México, México.; ABACUs-Centro de Matemáticas Aplicadas y Cómputo de Alto Rendimiento, Departamento de Matemáticas, Centro de Investigación y de Estudios Avanzados CINVESTAV-IPN, Carretera México-Toluca Km 38.5, La Marquesa, Ocoyoacac, Estado de México, 52740 México.; DIMNP UMR CNRS 5235, Université de Montpellier. France.; Centro de Ciencias de la Complejidad, Universidad Nacional Autónoma de México, México.; Instituto de Investigaciones Biomédicas, Universidad Nacional Autónoma de México, México.

## Abstract

Molecular regulation was initially assumed to follow both a unidirectional and a hierarchical organization forming pathways. Regulatory processes, however, form highly interlinked networks with non-hierarchical and non-unidirectional structures that contain statistically overrepresented circuits (motifs). Here, we analyze the behavior of pathways containing non-hierarchical and non-unidirectional interactions that create motifs. In comparison with unidirectional and hierarchical pathways, our pathways have a high diversity of behaviors, characterized by the size and number of attractors. Motifs have been studied individually showing that feedback circuit motifs regulate the number and size of attractors. It is less clear what happens in molecular networks that usually contain multiple feedbacks. Here, we find that the way feedback circuits couple to each other (i.e., the combination of the functionalities of feedback circuits) regulate both the precise number and size of the attractors. We show that the different sets of expected results of epistasis analysis (a method to infer regulatory interactions) are produced by many non-hierarchical and non-unidirectional structures. Thus, these structures cannot be correctly inferred by epistasis analysis. Finally, we show that the structures producing the epistasis results have remarkably similar sets of combinations of functionalities, that combined with other network properties could greatly improve epistasis analysis.

## Introduction

Early approaches considered that molecular regulation is composed of hierarchical and unidirectional interactions, where “above” molecules regulate “below” molecules, but molecules are not regulated by molecules at the same or lower levels (molecules usually represent genes and gene products^1^). Unidirectional and hierarchical interactions form *pathways,* comprised by an input, internal molecules, and an output (Fig.1A). Pathway dynamics (i.e., how the components in the pathway are activated and inhibited in time), follow a sequential order of regulatory events going from the input to the output through the internal molecules. Even though pathway dynamics seem to be an inherent property of molecular regulation^2, 3^, molecular regulation is usually complex, forming *highly inter-connected networks* that are neither unidirectional nor hierarchical^4–6^. Our objective is to systematically study the effect of including non-hierarchical and non-unidirectional interactions within pathways.

**Figure 1.**
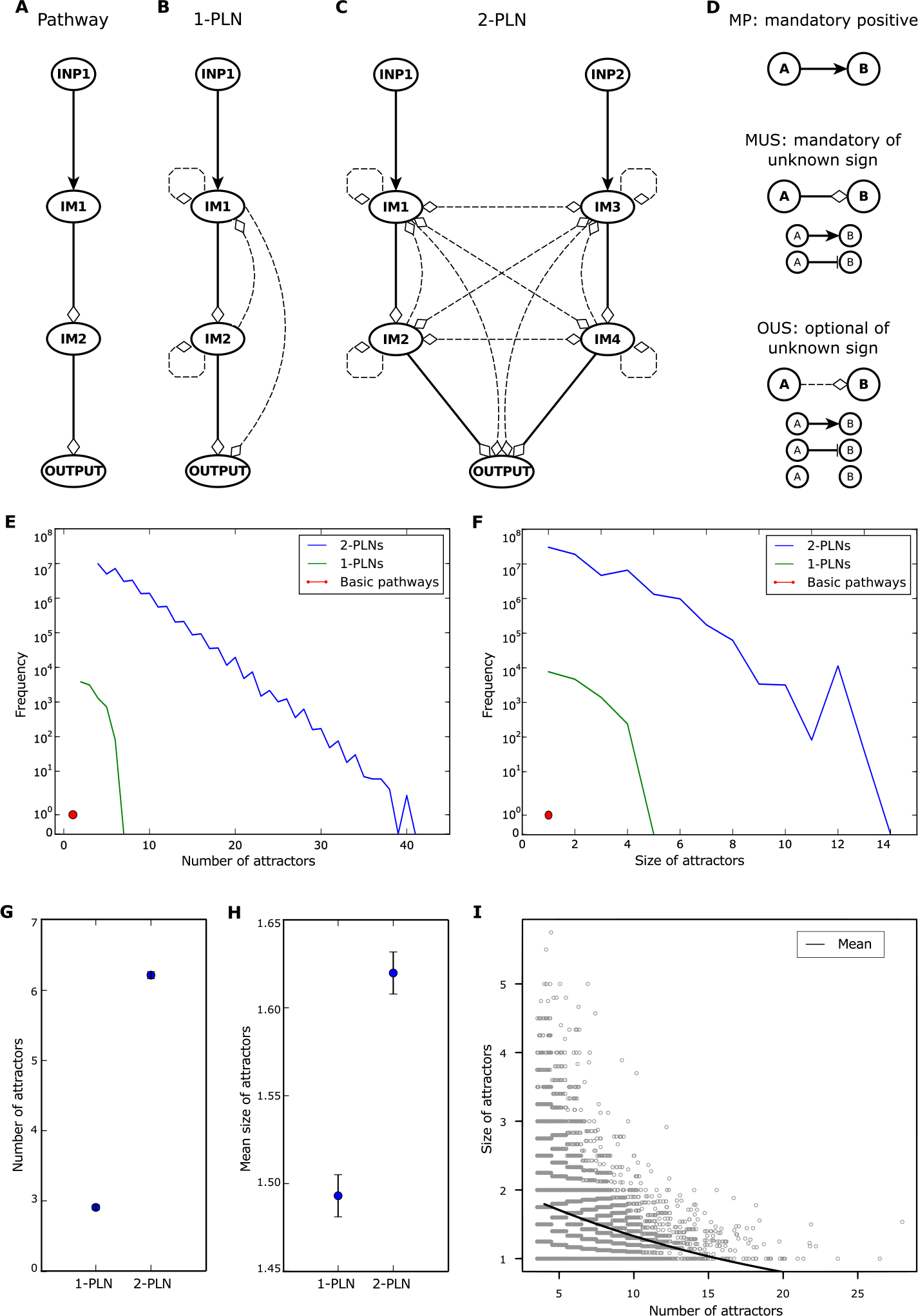
Pathway-like networks and their dynamical diversity. (A) Pathway, (B) 1-PLN and (C) 2-PLN structure. INP = INPUT and IM = Intermediary Molecules. (D) Interaction types considered in this work to construct pathways, 1-PLNs and 2-PLNs: mandatory positive (MP), mandatory of unknown sign (MUS) and optional of unknown sign (OUS). Mandatory interactions are always present, while optional in OUS interactions can be either present or not. Positive interactions are always positive. Unknown sign interactions can be either positive or negative. The possible combinations of negative and positive interactions of (A) form the four pathway variants, shown in Fig. 4B. Distribution of the (E) number and (F) size of the attractors. The red dot represents the result of the pathway, the green line the results of 1-PLNs and the blue line 2-PLNs. Mean and confidence interval of the (G) number and (H) size of attractors for 1-PLN and 2-PLNs. (I) Relation between the number and mean size of attractors of 2-PLNs. Each point represents a single 2-PLN data, while the line represents the values predicted by Poisson GLM. Points are displaced in the X axis only for visual purpose.

Adding interactions within a pathway modifies the pathway structure (i.e., the interaction graph describing who regulates whom), allowing for the appearance of regulatory motifs. Motifs are statistically overrepresented regulatory interactions found in molecular networks^5^. Among the motifs, circular chains of oriented interactions known as feedback circuits, are specially relevant, because they regulate both the number and size of the attractors. Attractors are stationary network states that represent biologically meaningful properties^7^, such as cell identity^8, 9^. Positive feedback circuits are necessary to have multiple attractors and negative feedback circuits are necessary to produce cyclic attractors^10, 11^. Moreover, the maximum possible number of attractors is regulated by positive feedback circuits^12^. It is not completely clear, however, how the precise size and number of attractors are regulated within molecular networks where many feedback circuits are present^13–15^. Here, we study how feedback circuits couple and regulate together the size and number of attractors.

Experimental research at small scales commonly uses traditional analyses that rely on the hierarchical and unidirectional assumptions. For example, epistasis analysis, as proposed by Bateson, is an analysis regularly used to infer and organize molecular regulatory interactions that assumes that regulatory interactions are hierarchical and unidirectional and can distinguish between different pathway structures^1, 16^. The presence of non-unidirectional and non-hierarchical interactions, however, can produce incomplete or even wrong gene regulation inferences when using epistasis analysis^17^. Thus, the study of the properties of non-hierarchical and non-unidirectional networks is fundamental to detect and solve limitations of traditional analyses. Here, we explore the capacity of epistasis analysis to infer pathways with non-hierarchical and non-unidirectional interactions and look for useful traits to distinguish between non-hierarchical and non-unidirectional regulatory structures.

In this work, we use synchronous Boolean networks to do a comprehensive analysis of Pathway-like networks (PLNs) that are non-hierarchical and non-unidirectional versions of pathways. We focus on the dynamical properties of PLNs, represented by both the number and the size of attractors. We show that PLNs have a large dynamical diversity in comparison with pathways. We confirm that feedback circuits are important regulators of the size and the number of attractors in a network. Then, we show for the first time, as far as we know, that the exact size and number of attractors in networks with multiple feedback circuits is in large part determined by the combination of the functionalities of feedback circuits. This combination refers to the network states where the feedback circuits in a network have an actual effect over the network dynamics. Then, we study the effect of adding non-unidirectional and non-hierarchical interactions on the reach of epistasis analysis to infer regulatory interactions. Epistasis analysis has different sets of expected results for each possible pathway structure. We show that there are a vast number of PLNs capable of producing each set of expected results. Thus, epistasis analysis produces incomplete or even wrong inferences when dealing with non-hierarchical and non-unidirectional regulatory structures. Interestingly, the epistasis analysis’ sets of expected attractors have the same number of attractors of the same size and are produced by PLNs with remarkably similar sets of combinations of functionalities. Thus, we explore how to use the combination of functionalities combined with both the network dynamic and structure to improve epistasis analysis.

## Methods

### Boolean networks

Molecular networks have variables representing the genes, mRNA, proteins, or any other type of molecules included in the network. The network structure is represented by the network’s interaction graph, which is a directed graph that describes the regulatory interactions between the network’s variables. In this work, we use the Boolean formalism to model molecular networks for the following reasons. (1) Because of their simplicity, Boolean networks are well suited to perform analyses in a large number of networks, without needing to deal with problems such as parameter estimation. (2) Despite their simplicity, Boolean networks obtain biologically meaningful results (e.g.,^8, 9, 18–20)^. (3) Feedback circuits are fundamental for this work, and feedback circuit functionality is well studied in Boolean networks^21, 22^. (4) Here we will focus on the size and number of attractors, and it has been proven that in Boolean networks positive and negative feedback circuits are a necessary condition for multistability and oscillations, respectively ^11, 23^. (5) We will use epistasis analysis that assumes that the molecules behave as Boolean variables^1, 16^. Hence Boolean networks are a natural and simple extension of epistasis analysis.

In Boolean networks, variables can only take one of two possible values, 0 or 1, and their dynamics is described by

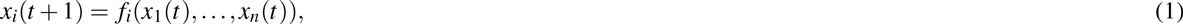

where *x_i_(t+1)* represents the value of variable *i* at the time *t+1* as a Boolean function *f_i_* of its *n* regulators *x_1_(t),…,x_n_(t)* at the current time. In particular, we use synchronous Boolean networks, where the value of all variables is updated at each time step.

The set of all variables values at time *t* is a network state. Stationary network states are known as attractors. Single-state, stationary configurations are known as fixed-point attractors whereas a set of network states that orderly repeat correspond to cyclic attractors. The size of an attractor is equal to the number of network states that conform such attractor.

For this work it is important to note that not all interactions in a network structure are necessarily functional. Intuitively, an interaction from a variable *i* to a variable *j* is considered *functional,* if *j* can change its value due to a change only in the value of i. The interaction sign is positive, if the change in the value of *j* goes in the same direction as the change in the value of *i*, and is negative otherwise (see Supplementary information for all the formal definitions). Note that a variable can act as a positive regulator and as a negative regulator of the same variable in different states. Interactions where a variable is a positive and negative regulator of another variable are not common in molecular networks and non-functional interactions do not provide interpretable results. Hence, in the networks analyzed here, we forbid both non-functional interactions and interactions where a regulatory variable has both a positive and a negative influence over another variable.

A feedback circuit is a set of directed interactions forming a closed path. Feedback circuits can be positive or negative. The sign of a feedback circuit is given by the signs of its interactions. A circuit is positive if it has an even number of negative interactions, it is negative otherwise. Similarly to the interactions, the sole presence of a circuit in a network does not guarantee its functionality. Thus, a circuit is considered functional if all interactions of the circuit are functional in a set of common network states^21^. The functionality of a feedback circuit is characterized by the sign and number of network states where the circuit is functional. The combination of the functionalities of feedback circuits is the set of all circuits functionalities present in a given network. As a network can have many circuits and each circuit can have many functionalities, a network structure could produce many different combinations of functionalities.

### Pathway-like networks construction

We analyze two types of networks, namely pathways and pathway-like networks (PLNs), both containing different types of interactions identified with the acronyms MUS, OUS and MP, which stand for mandatory unknown sign, optional unknown sign, and mandatory positive, respectively (Fig.1D). MUS and MP interactions form a unidirectional and hierarchical pathway structure (Fig.1A). PLNs contain MUS, MP and OUS interactions (Fig.1B and C). We construct two PLNs variants: single (1-PLNs) and double (2-PLNs). 1-PLNs is the set of networks that contains a pathway within its structure and at least one OUS interaction (Fig. 1B). 2-PLNs is the set of networks composed by two parallel pathways regulating the same output and at least one OUS interaction (Fig. 1C). 2-PLNs may have cross-regulation among their constituent pathways, which is a common biological situation. The classical pathway structure used in the epistasis analysis is shown in Fig.1A. Note that the added interactions in 1-PLNs and 2-PLNs create motifs, including feedback circuits.

We used two different approximations to analyze the size and the number of attractors. (1) We simulated each PLN starting from all its initial conditions until finding all the attractors. The number of initial conditions in any network is equal to 2^*v*^, where *v* is the number of variables. (2) We use symbolic algorithms to search for PLNs with a specific number and size of attractors. Is important to note that in most cases, a given network structure can be described by more than one set of Boolean functions. Thus, there is a vast number possible PLNs structural and dynamical variants. Without considering the sign of the interactions, the number of possible structures for a network with *v* variables is equal to 2^*v*^2^^. The Boolean functions associated with a variable is 2^2^*r*^^, where *r* is the number of regulators of the variable. Therefore, the total number of possible Boolean functions for a completely interconnected network with *v* variables is 2^n×2^n^^. The analysis of such a number of variants quickly becomes unfeasible. In particular, there are ≈ 4.15 × 10^34^ 2-PLNs. Thus, we use random sampling or constrained the maximum number of interactions for 2-PLNs analyses (see Supplementary information for more details).

## Results

### Non-hierarchical and non-unidirectional interactions greatly increases the dynamical diversity of pathways

Conventionally, molecular regulation is represented as unidirectional and hierarchical pathways, comprised by an input, internal molecules, and an output (Fig. 1A). To study pathways with more realistic structures, we consider the possibility that the internal molecules may regulate any component inside the pathway (excluding the input). We call this structure a *Pathway-like network* (PLN). We analyzed single (1-PLNs) and double (2-PLNs) PLNs (Fig. 1B and C). PLNs have a non-unidirectional and non-hierarchical structure. Regarding their dynamics, PLNs are constructed in such a way that the generation of Boolean functions producing “meaningless” behaviors is forbidden^17, 24^. Altogether, PLNs contain realistic structural and dynamical properties (see Methods).

We analyzed the dynamical diversity, measured as the number and the size of attractors in pathways, 1-PLNs and 2-PLNs. We observed that the dynamical diversity vastly increases from pathways to 1-PLNs to 2-PLNs (Fig.1 E and F). Because the inputs follow the identity function, the minimum number of attractors is equal to 2^|*inputs*|^, where *inputs* is the set of inputs. The maximum number of attractors, on the other hand, are 2, 6, and 44 in pathways, 1-PLNs and 2-PLNs, respectively. This increase is also observed in the mean values, where the mean value of the number of attractors of 2-PLNs is significantly larger than in 1-PLNs (*P* < 0.001; Fig.1G; see Supplementary information file for detailed information about all statistical results). Similar results can be observed for the size of attractors. Specifically, the maximum sizes are 1, 4, and 13 for pathways, 1-PLNs and 2- PLNs, respectively. Here, too, the mean value of the size of the 2-PLNs attractors is significantly larger (*P* < 0.001; Fig.1H). Also, there is a statistically significant negative relation between the mean size and mean number of attractors (*P* < 0.001; Fig. 1I and Supp. Fig.1A). Note that both the mean size and number of attractors are much closer to their minimum value than to their maximum value. This indicates that most networks have a small number of attractors of small size; and at first sight it seems that they might fit a long-tailed distribution. It is interesting to note that these data do not fit power-law, logarithmic, exponential, normal or Poisson distributions. Nevertheless, our results clearly show that the overall diversity of dynamical behaviors grows from pathways to 1-PLNs to 2-PLNs, represented by the number and the size of attractors.

### Feedback circuits role in the regulation of the PLNs attractors properties

To understand what regulates the observed increase in the number and the size of attractors, we focus on feedback circuit motifs. Positive feedback circuits are necessary for multistability, while negative feedback circuits are necessary to produce cyclic attractors^10, 11^. Thus, it seems reasonable to look for relations between the number and size of attractors with the presence of positive and negative feedback circuits. Indeed, positive feedback circuits are positively related with the number of attractors (*P* < 0.001), and negatively related with the size of attractors (*P* < 0.001). In turn, negative feedback circuits have a negative relation with the number of attractors (*P* < 0.001), and a positive relation with the size of attractors (*P* < 0.001; Fig. 2A-D; Supp. Fig.1).

**Figure. 2.**
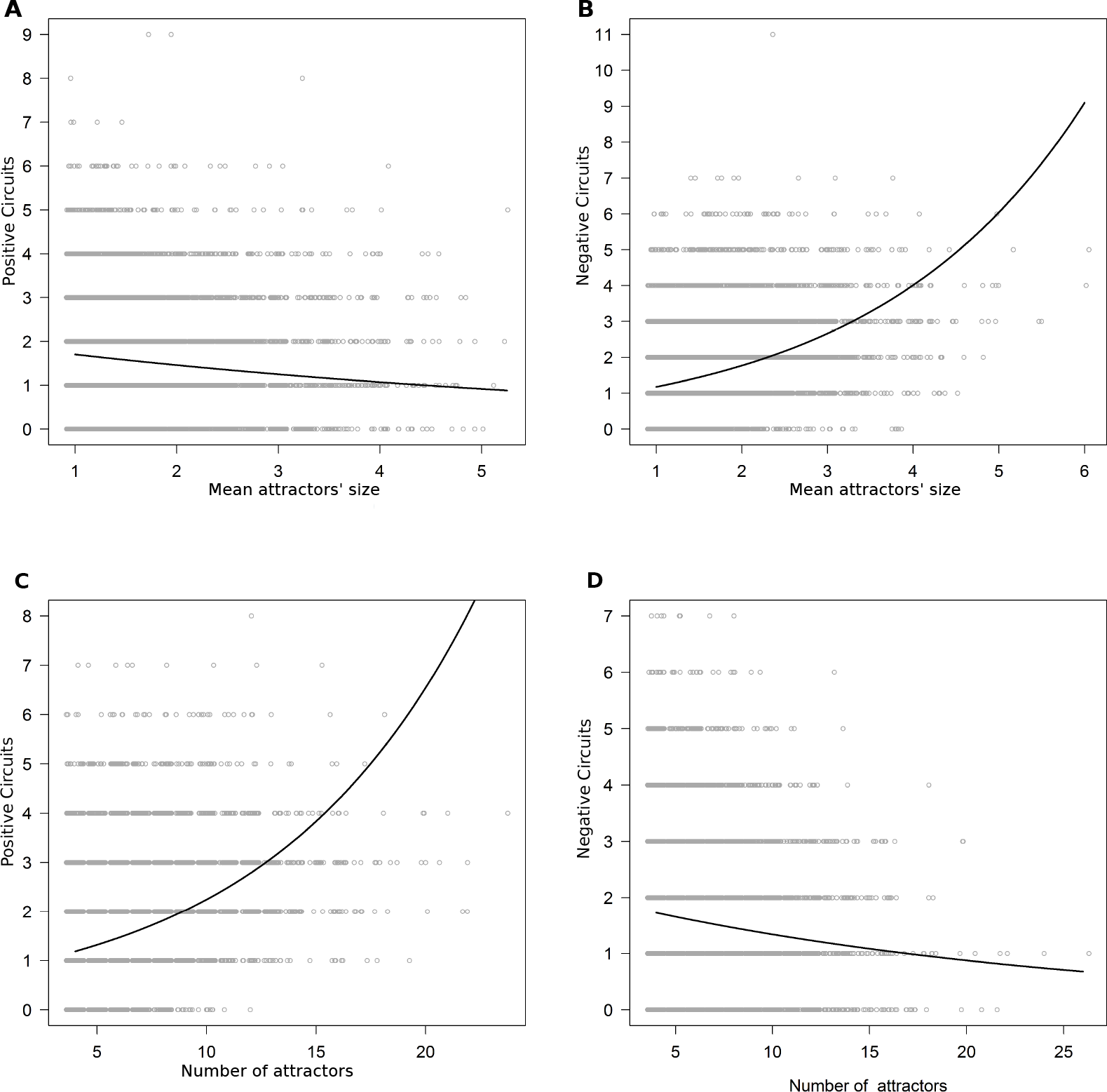
Relationship between 2-PLNs circuits and attractors. Mean size of attractors vs. quantity of (A) positive and (B) negative feedback circuits, respectively. Number of attractors vs. quantity of (C) positive and (D) negative feedback circuits, respectively. The lines are predicted by Poisson GLMs. Points are displaced in the X axis only for visual purpose. Similar results were found in 1-PLNs (Supp.Fig. 1B-E).

To better analyze the effect of feedback circuits on the attractors, we classified the PLNs according to their number and size of attractors in four categories. (1) PLNs with the minimum number of attractors, whose mean size were bigger than 90% of the mean of the total set of PLNs analyzed (n-s+). (2) PLNs with more attractors than the 90% of PLNs, all fixed-point attractors (n+s-). (3) PLNs with the minimum number attractors, all fixed-point (n-s-). (4) PLNs with more attractors than 90% of the PLNs, whose size was bigger than 90% of the PLNs (n+s+) (Fig.3A). Then, to see if the quantity of feedback circuits in a PLN is important to determine the number and the size of the attractors, first, we compare the total number of feedback circuits of n+s+, n+s-, n-s+ and n-s- PLNs against all the PLNs analyzed. The total number of circuits, does not differs between any selection of 1-PLNs and the complete set of 1-PLNs, but it differs between all possible 2-PLNs selections and the total set of 2-PLNs (Fig. 3B and C and Table 1). Then, we analyzed both the number of positive and the number of negative feedback circuits. In comparison with all 2-PLNs analyzed, 2-PLNs n+s- and n-s- have significantly fewer positive and fewer negative feedback circuits, while 2-PLNs n-s+ and n+s+ have significantly more positive and more negative feedback circuits. In 1-PLNs n+s- and n-s- there are significantly more positive and fewer negative feedback circuits, while the opposite behavior is observed in 1-PLNs n-s+ (Table 1). As observed, the results differ from 1-PLNs to 2-PLNs, and it is not possible to understand how the size and the number of attractors is defined from the number of feedback circuits, whether it is the total, the positive or the negative feedback circuits.

**Figure 3.**
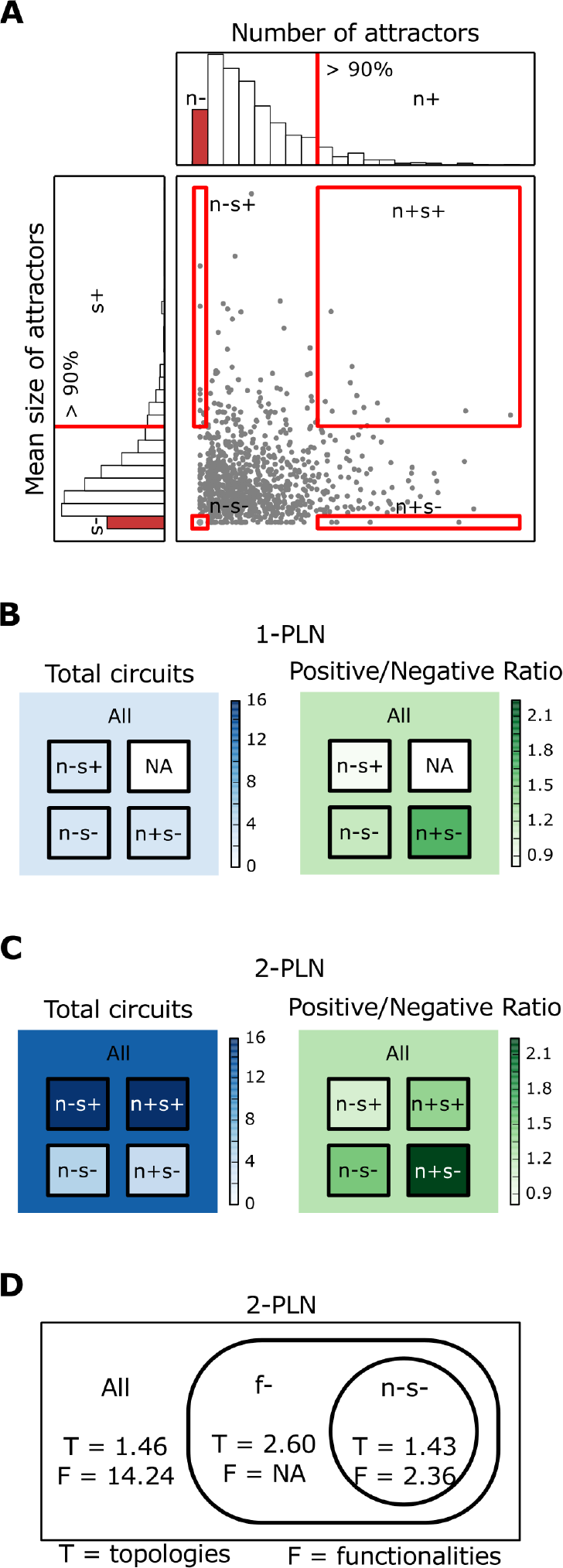
PLNs attractors properties regulation by feedback circuits. (A) Classification of PLNs according to the number and mean size of the attractors: *n*+ more attractors than 90% of the PLNs, *n*- minimum number of attractors, *s*+ mean size of attractors bigger than 90% of the PLNs and *s*- fixed-point attractors. Mean number of total feedback circuits and positive/negative feedback circuits ratio for all PLNs and for each of the possible classifications: (*n*-*s*-, *n*+*s*-, *n*-s+, *n*+*s*+) in 1-PLNs (B) and 2-PLNs (C). (D) Mean number of combinations of functionalities in a structure (F) and mean number of PLNs structures that contained each combination of functionalies (T) in all 2-PLNs (All), 2-PLNs *n*-*s*- (n-s-) and all 2-PLNs that contain the *n*-*s*- combinations of functionalities (f-). NA stands for non applicable.

**Table 1.**
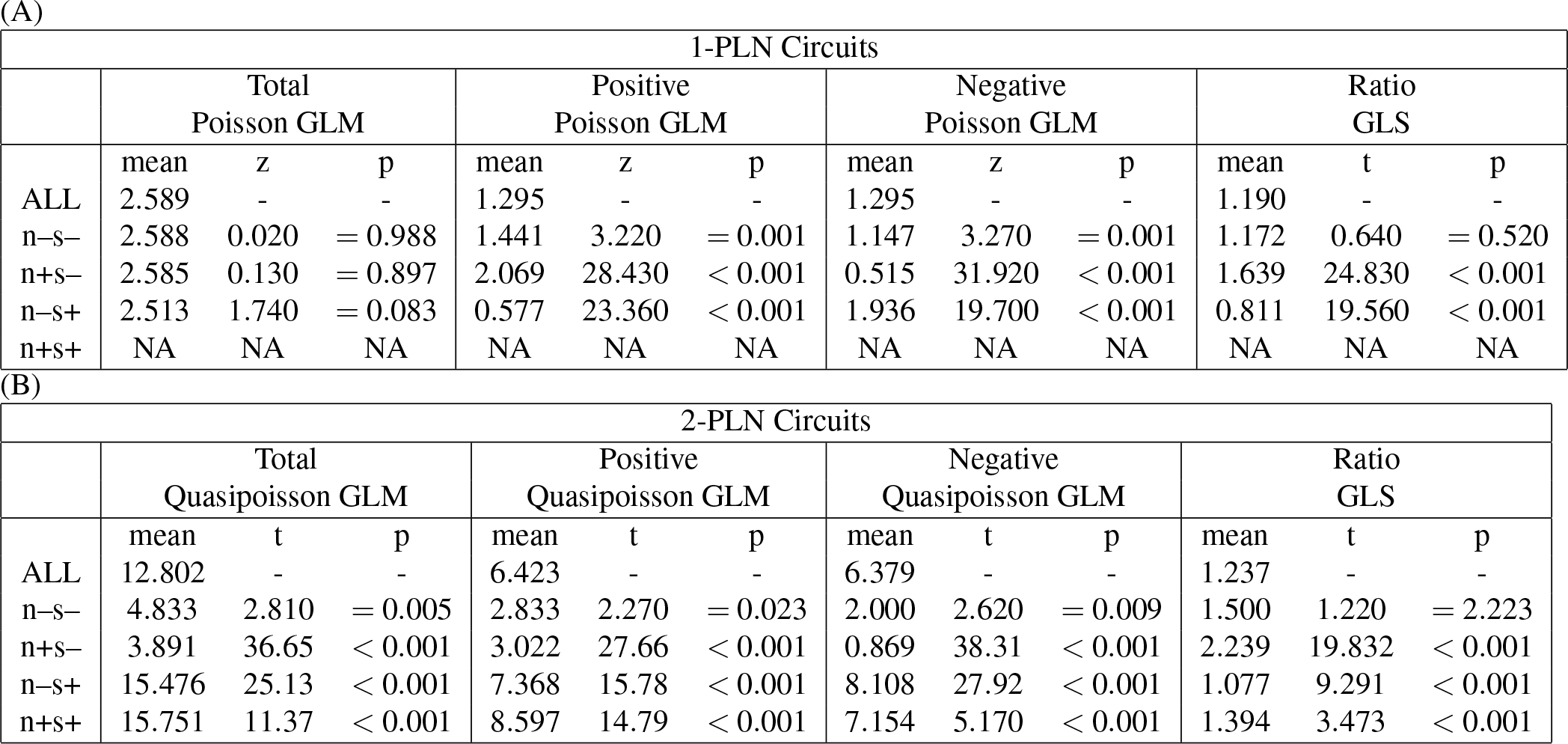
Feedback circuits analyses. Total, positive, negative and positive/negative ratio of feedback circuits in (a) 1-PLNs and 2-PLNs (B). All values are from comparing ALL against each of the PLNs selection (i.e., n-s-, n-s+,n+s- and n+s+).

According to Kwon et al.^25^, the positive/negative feedback circuits ratio provides the trend of the number and the size of attractors, with larger ratio values for networks with more attractors of smaller size. If this is the case, n+s- PLNs should have a bigger positive/negative feedback circuits ratio than the complete set of PLNs. On the other hand, n-s+ should have a lower positive/negative feedback circuit ratio than the complete set of PLNs. Finally, n-s- and n+s+ could have a balance between positive and negative feedback circuits and not show any difference in their ratio against the complete set of PLNs analyzed. As observed in Fig.3B and C and Table 1, this is indeed the case in both 1-PLNs and 2-PLNs (*P* < 0.001). From the circuits ratio result, we can expect the PLNs n+s- to have more positive feedbacks and fewer negative feedbacks, the opposite behavior for n-s+, and a similar number of positive and negative feedback circuits in the n+s+ and n-s-. As observed in Table 1, this is the case in both 1-PLNs or 2-PLNs. Thus, our results support the idea that the positive/negative feedback circuits ratio provides the correct trend for the size and the number of attractors.

### The combination of the functionalities of the feedback circuits regulates the attractors size and number

We detected PLNs n+s- with a lower positive/negative feedback circuits ratio than PLNs n-s+ and vice versa. This result is opposite to what is expected. The positive/negative feedback circuits ratio cannot explain this type of results, as it do not provides a mechanistic of how do multiple feedback circuits regulate the size and the number of attractors. Is important to note that the mere presence of a feedback circuit in the structure of a network does not guarantee the expected behavior unless the circuit is functional^21, 26^. For a circuit to be functional, all interactions within such a circuit should be functional at the same time^21^. The functionality of the circuits could explain why we can find larger positive/negative feedback circuits ratio in n-s+ than in n+s- and vice versa. For example, a network with the same number of positive and negative feedback circuits, but whose negative feedback circuits are not functional, could produce a large number of small attractors, even when the positive/negative feedback circuits ratios is equal to 1. We find that at least one feedback circuit is functional in all the PLNs with feedback circuits in its structure. However, not all feedback circuits are functional. This indicates that, (1) due to the constraints imposed to the PLNs Boolean functions (see Methods), at least one feedback circuit within a PLN will be functional, but (2) the functionality of each feedback circuit within a PLNs with multiple feedback circuits will depend on how it interacts with the other feedback circuits and (3) that the coupling among feedback circuits could be behind the regulation of the attractors size and number. Observe that because the definition of circuit functionality does no imply that the circuit should be functional in a specific set of network states, it is possible to have a set of functionalities for the same feedback circuit in a given network structure. Furthermore, since a network can have more than one feedback circuit, it is also possible for a single network structure to have different *combinations* of the functionalities of its feedback circuits (hereinafter named combinations of functionalities). Different combinations of functionalities reflect different couplings between feedback circuits. Consequently, we analyzed the combination of functionalities to see if they regulate the size and number of attractors.

To study if the combinations of functionalities are the main regulators of the attractors size and number, we analyzed such combinations in PLNs containing at least one feedback circuit. In total, all 1-PLNs (8,960) and a sampling of one million 2-PLNs were analyzed. From these PLNs, we selected the n-s-PLNs to compare them with the total of PLNs. We selected n-s- PLNs for this analysis, because they are the more common ones, providing the possibility of finding each combinations of functionalities in more than one PLN in both 1-PLNs and 2-PLNs. We found 680 and 61,951 n-s- 1-PLNs and 2-PLNs, respectively. In the set of all PLNs, we found 62 and 231,949 combinations of functionalities for 1-PLNs and 2-PLNs, respectively. In the PLNs n-s- there were 8 and 11,260 combinations of functionalities.

We noticed that the number of PLN structures was reduced from all PLNs to PLNs n-s- from 14 to 6 and from 23,786 to 6,983 in 1-PLNs and 2-PLNs, respectively. Given that a combination of functionalities can only be achieved when the PLN structure contains the feedback circuits that conform with such a combination, we performed an analysis to assure that the reduction in the combinations of functionalities number is not due to the reduction in PLNs structures. If the PLN structures reduction are the main cause in the reduction of the number of combinations of functionalities, we should find similar sets of combinations of functionalities in a structure, independently of whether we analyze the complete set of PLNs or the n-s- PLNs. We found that the mean number of combinations of functionalities contained in each structure diminished from 8.85±21.38 to 2.66±3.65 and from 14.24±0.199 to 2.36±0.146 in 1-PLNs and 2-PLNs (Fig.3D), respectively. This reduction is significant in the 2-PLNs case (*P* < 0.001). These result indicates that even when the PLNs structure determines the combinations of functionalities that can be generated, only some of such combinations of functionalities are able to produce n-s- PLNs, suggesting that the combinations of functionalities is an important regulator of the attractors size and number.

If the combinations of functionalities are the main regulator of the attractors size and number, any or almost any PLN capable of producing the same combinations of functionalities should also be able to produce the same number and size of attractors, independently of the PLN structure. In the case of the 1-PLNs, each combination of functionalities is contained in exactly two structures in both the n-s- 1-PLNs and the complete set of PLNs. In 2-PLNs there was no significant difference (*P* = 0.331) as, on average, each combinations of functionalities is present in 1.46±0.005 in the complete set of 2-PLNs, vs. 1.43±0.0022 in the 2-PLNs n-s- (Fig.3D). Hence, our results strongly suggest that the combinations of functionalities are a fundamental regulator of the PLN’s attractors properties.

Finally, to study if there were other important constrains in the PLNs structures for the appearance of n-s- attractor properties besides the combination of functionalities, we analyzed which structures in the complete set of PLNs contained the functionalities found in the PLNs n-s-. We found that each combination of functionalities found in the n-s- PLNs was contained in 2±0 and 2.60±0.030 in the complete set of 1-PLNs and the 2-PLNs, respectively. As observed, there is no change in the 1-PLNs. However, there is a slight, but significant increase in the 2-PLNs case (*P* < 0.001) (Fig.3D). Hence, we looked if this increase was due to the need of another interaction (or a lack of it) to produce the desired results. We found no indispensable additional interactions. Hence, the increase should be due to another type of constrains that we were not able to detect here. Our results indicate, however, that the combinations of functionalities are one of the most important regulators of the attractors properties.

### Combinations of the functionalities of feedback circuits and the analysis of epistasis

Our results suggest that, in general, networks with the same combination of functionalities produce the same number and size of attractors. It is important to note, that this result seems to be independent of which network states represent the attractors, as we did not considered this data in our previous analyses. Thus, a complementary way to test if the combinations of functionalities regulate the number and the size of attractors, is to search for networks with the same number of attractors of the same size, but whose attractors are represented by different network states. These networks should be produced by the same or similar sets of combinations of functionalities. Observe that the possible combinations of positive and negative interactions in a pathway already gives four pathway variants that have the same number and size of attractors, but the attractors of each pathway variant correspond to different network states. The epistasis analysis, as originally conceived by Bateson, has characterized the expected results of the four pathway variant^1, 16^. Thus, we used the expected results of each pathway variant by the epistasis analysis to further analyze the importance of the combinations of functionalities in the regulation of the size and number of attractors.

Briefly, epistasis is a term used when the phenotype of an allele is masked by an allele in another locus. The gene with the allele whose phenotype persists when the alleles of both loci are present is called epistatic gene, while the other is the hypostatic gene. Epistasis analysis uses a simple set of two rules to order the epistatic and hypostatic genes^1, 16, 27^. First, in a double-mutant experiment, the epistatic gene is upstream and positively regulates the downstream gene when the two genes used in the double-mutant display a characteristic single-mutant phenotype under the same condition. And second, in a double mutant experiment, the epistatic gene is downstream and is negatively regulated by the upstream gene when the two genes display a characteristic single mutant phenotype under different conditions. The epistasis analysis can be formalised in Boolean term in a straightforward way (Fig.4A). We named each pathway variant according to the sign of the regulation from GENE1 to GENE2 and from GENE2 to the OUTPUT, as ++, +-, -+ and - -. Then, we interpreted the epistasis analysis expected results for each pathway variant as the set of target attractors in our PLNs (Fig 4B) and analyzed the number of 1-PLNs and 2-PLNs that achieved each of the fours possible sets of target attractors.

**Figure 4.**
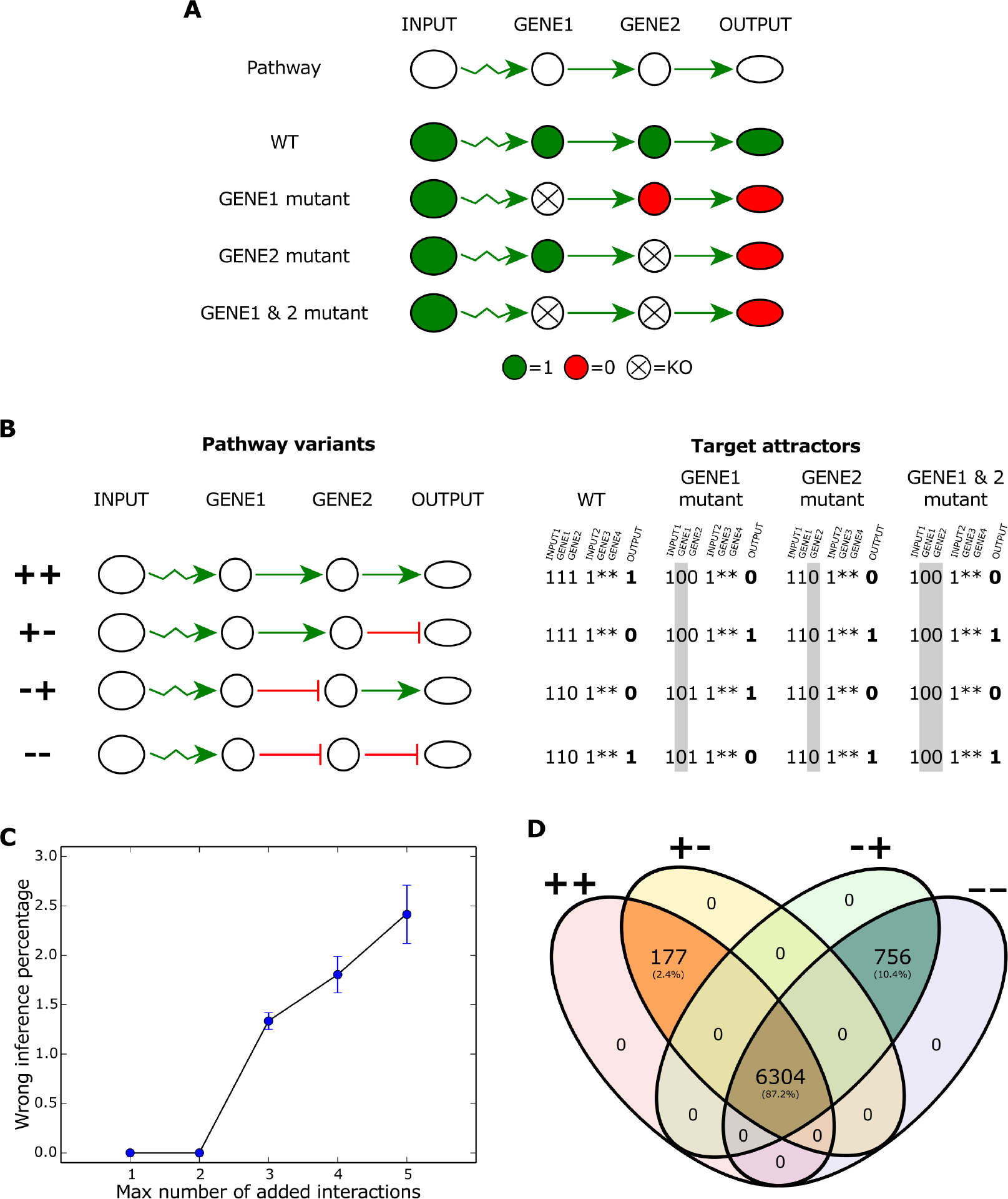
Characterization of the PLNs that produce the epistasis results. (A) Boolean interpretation of the epistasis analysis using the ++ pathway variant shown in (B). Target attractors for each of the four pathway variants using the epistasis analysis. Mutated genes are highlighted in grey. For 1-PLNs we only consider INP1, GEN1 GEN2 and OUTPUT values. Epistasis analysis do not consider the possibility of a second parallel and cross-talking pathway. Thus, the asterisks represent unknown values for GENE3 and GENE4 in the 2-PLNs case. Input values are considered equal to 1 because the epistasis analysis assumes that the input values are constant during the experiment^1^. (C) Percentage of 2-PLNs that produce the target attractors of different pathway variant than the pathway variant contained in the 2-PLN as we added more interactions the unidirectional and hierarchical pathway structure. (D) Percentage of combinations of functionalities shared between the PLNs that produced each of the four sets of target attractors.

The number of PLNs able to produce the target attractors in the 1-PLNs is 60 for the ++ and +- variants and 68 for the -+ and - - variants. We were unable to perform an exhaustive 2-PLNs search because of the astronomical number of redundant 2- PLNs found. We restricted our search to 2-PLNs with a maximum of five extra interactions compared to the pathway structure. We found than more than 4.731 × 10^7^ 2-PLNs that produced each set of target attractors.

In the 1-PLNs case, the target attractors of a pathway variant are only obtained when the same pathway variant is contained in the PLN. On the other hand, ≈ 2.41% of the 2-PLNs produced the target attractors of a pathway variant different from the pathway variant contained in its structure. This demonstrates that epistasis analysis can produce wrong regulatory inferences when there analyzing non-unidirectional and non-hierarchical networks, as stated before by our group^17^. The wrong inferences increase as we added more interactions (Fig 4C) and are produced thanks to the added interactions between the parallel pathways contained in the 2-PLNs. It is interesting to note, that when there are wrong inferences, the extra interactions produce an alternative pathway that conformed with the expected pathway variant but that contained some intermediary steeps between GEN1 and GEN2 or between GEN2 and OUTPUT (see some examples in Supp.Fig. 2). However, the alternative pathway by itself is not sufficient to produce the expected results as the alternative pathway can be created by adding one interaction and inconsistencies between the target attractors and the pathway variant contained in the PLNs appear only when we added three or more interactions (Fig 4C). Thus, our results demonstrate that epistasis analysis produce wrong and incomplete inferences in non-hierarchical and non-unidirectional networks, but when a network has a low connectivity, wrong regulatory inferences are scarce.

The combinations of functionalities that produced the four target attractors are almost the same in both 1-PLNs and 2-PLNs. In 1-PLNs all four target attractors share one combination of functionalities. This is a meaningful result as that is the only combination of functionalities that produced the target attractors of the variants ++ and +- and one of the two combination of functionalities that produce the target attractors of the variants -+ and - -. In the 2-PLNs case, there are 6,481 combinations of functionalities that produced the target attractors of the variants ++ and +- and 7,060 that produced the target attractors of the variants -+ and - -. 6,304 combinations of functionalities found are shared (Fig.4D). This is an astonishing result that greatly supports the importance of the combinations of functionalities to determine the attractors properties, as the target attractors vary for each pathway variant, indicating that the common feature among the PLNs found are the number and the size of attractors.

### Comprehensive characterization of PLNs

Under the unidirectional and hierarchical assumptions, there is only one structure capable of producing the attractors expected in each pathway variant. However, our results show that adding non-hierarchical and non-unidirectional interactions in a pathway structure increases the number of networks that can produce these attractors. This raises a problem for the analyses of molecular regulation by traditional methods, such as the epistasis analysis, as they cannot distinguish between these networks. Hence, we did an exploration of which properties could be useful to distinguish between these networks. For this characterization, we used all 1-PLNs and 2-PLNs with no more than 2 interactions added to the unidirectional and hierarchical pathway structure, comparison of PLNs with three or more extra interaction was extremely challenging or computationally impossible.

First, we analyzed if structurally similar PLNs followed similar dynamics. As observed in fig.5A, structurally close PLNs can have big dynamical distances and vice versa, indicating that dynamical and structural distances are not related. On the other hand, we noticed that PLNs with the same combination of functionalities and the same structure cluster together and that the PLNs within the same cluster have more similar dynamics between them, than PLNs in a different cluster (Fig.5B). Thus, each cluster have distinguishable properties. Consequently, the combination of functionalities, seem to group together dynamical and structural similar PLNs. Thus, studying the characteristics of these regions could be an important step towards a better and more general understanding of molecular regulation and could allow for a better characterization of regulatory networks to improve the scope of traditional methods.

**Figure 5.**
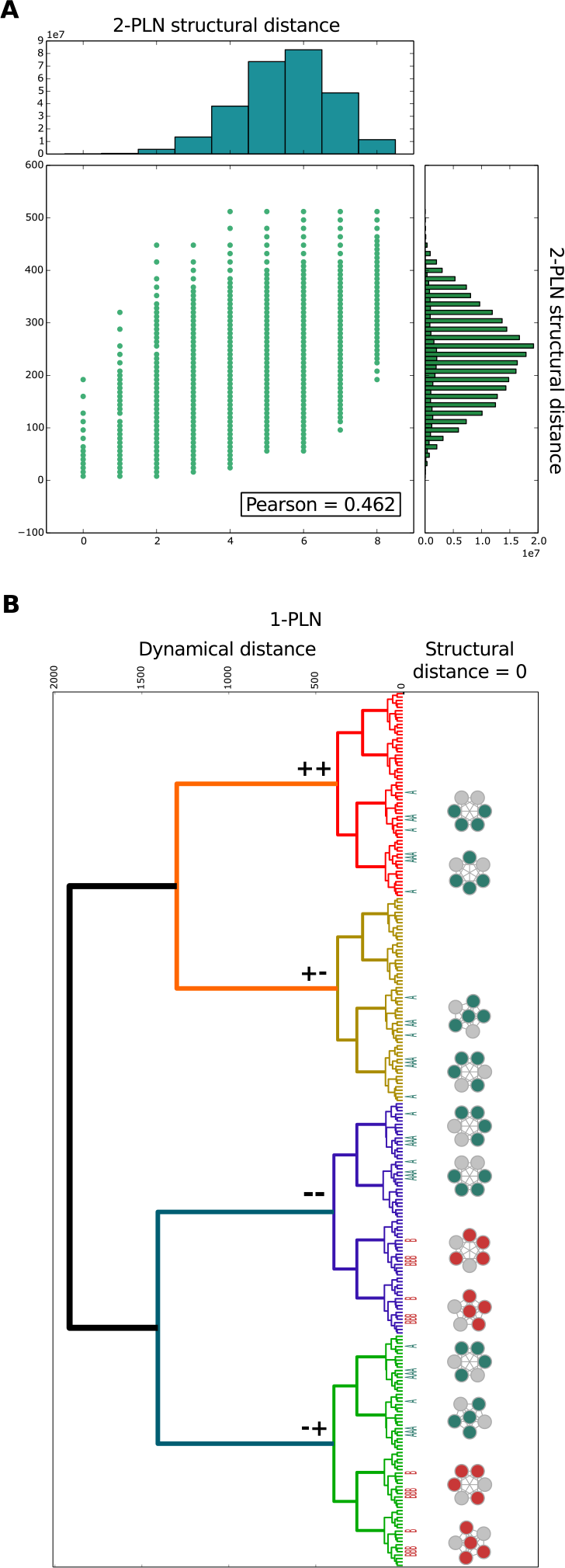
PLNs characterization. (A) Relationship between dynamical and structural distance between 2-PLNs. (B) In the right size, 1-PLNs hierarchical clustering dendogram using the dynamical distance. The 1-PLNs cluster according to the pathway variant. The colored leaves represent the combinations of functionalities observed on the left. On the left, 1-PLN cluster formed with structural distance of 0, the grey color correspond to PLNs with no functional circuits, and the other colors correspond to the different combinations of functionalities found. Similar results were obtained in 2-PLNs.

### Conclusions and discussion

We characterized the effect of adding certain interactions within unidirectional and hierarchical pathway structures. These extra interactions create realistic regulatory structures with a resulting non-unidirectional and non-hierarchical organization containing motifs that we named Pathway-like Networks (PLNs). Additionally, we included certain procedures to ensure the creation of only biologically meaningful dynamics^3, 18, 24^. As a result, PLNs have realistic structural and dynamical properties. Here we showed that PLNs have a great dynamical diversity, as characterized by the number and the size of attractors.

The explosion in the dynamical possibilities of the PLNs was expected from previous works. For example, Kauffman^7^ hypothesized that the number of attractors in random Boolean networks increase as 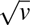. Later, it was found that the number of attractors in Boolean networks increases faster than any power law as the number of variables increases^28^. A strict comparison between these proposals and our results is difficult, because both Samuelsson and Kauffman’s proposals were based on studies with a fixed number of connections, while our networks have a variable connectivity. Nonetheless, our results clearly show that PLNs have more attractors than unidirectional and hierarchical pathways.

We then focused on the role of network motifs, specially feedback circuits, in the size and the number of attractors. We found that, as the number of positive feedback circuits increases, the number of attractors increases and the size of the attractors decreases. We also observed the exact opposite relation between the number of negative feedback circuits and the number and size of attractors. These results were expected considering that positive and negative feedback circuits are required for multistability and oscillations^10, 11^. As it has been reported^25^, the positive/negative feedback circuits ratio give the correct trend for the number and the size of the attractors. These results are interesting, but they do not explain how feedback circuits in PLNs, which can have multiple coupled feedback circuits, regulate the size and number of attractor. Thus, we looked for a more mechanistic understanding of the regulation of the size and the number of attractors by analyzing the coupling of feedback circuits. We found that the combination of functionalities (i.e., the way feedback circuits couple) is a key regulator of the number and the size of attractors. In general, PLNs with the same combination of functionalities can produce the same number and size of attractors, independently of its structure. In our opinion, this is a characteristic worth of further research to fully understand the nature of molecular regulation.

In accordance with previous observations^13, 19, 29^, we found that many PLNs are capable of producing the same number, size, and even the same set of attractors. This result emphasizes how limited are the traditional methods for the analysis of experimental results, as the epistasis analysis. First, because such analyses consider only a restricted number of possible networks, they are not well suited to deal with the huge diversity of possible dynamic behaviors. Second, because such methods are unable to distinguish between alternative networks producing the same set of attractors. Thus, we searched for the number of PLNs that produced the epistasis results. In 2-PLNs, the number of network that produced the same set attractors was so huge, that we needed to constrain our search to a limited quantity of PLNs structures. As a consequence, we can conclude that with the use of traditional methods for the analysis of experimental data many interactions cannot be detected^14^. Even more, we find that in some cases they can even produce wrong gene regulation inferences^17^. These incorrect inferences are due to the appearance of alternative pathways that can produce the expected behaviors. There may not be general rules to infer complex networks structure, such as PLNs^13, 14^. However, the PLNs found, share most of their combinations of functionalities and the PLNs with the same structure and with the same combination of functionalities produce dynamically similar regions that can be distinguished. Thus, we believe that a more general understanding of the combination of functionalities and its relation with networks structure and dynamic will open possible ways to study and analyze molecular regulation of biological processes.

## Acknowledgements

We thank Gustavo Ramos, Luis Cauich and Jocelyn Champagnon for statistical advice. We thank Dr. Emmanuel Faure, Dr. Mariana Benitez and Dr. Christophe Godin for their comments and discussion. We also gratefully acknowledge support form PASPA-DGAPA-UNAM and Conacyt grants 221341 and 261225. LM acknowledges the sabbatical scholarships from PASPA-DGAPA-UNAM and CONACYT 251420.

## Author contributions statement

EA, LM and ERAB conceived the project. EA and LM designed the research. EA, LM, NW, SM and MEMS performed the research. EA, DGT, MEMS, AN analyzed the data. EA, NW, LM, SM, AN and DR provided and developed software tools. EA, MEMS and DGT prepared the figures. EA and LM wrote the manuscript with inputs from all authors.

## Additional information

**Competing financial interests**: The authors declare no competing financial interests.

